# Genomic characterization of the antiviral arsenal of Actinobacteria

**DOI:** 10.1101/2023.03.30.534874

**Authors:** H. Georjon, F. Tesson, H. Shomar, A. Bernheim

## Abstract

Phages are ubiquitous in nature, and bacteria with very different genomics, metabolisms, and lifestyles are subjected to their predation. Yet, the defense systems that allow bacteria to resist their phages have rarely been explored experimentally outside a very limited number of model organisms. Actinobacteria are a phylum of GC-rich gram-positive bacteria, which often produce an important diversity of secondary metabolites. Despite being ubiquitous in a wide range of environments, from soil to fresh and sea water but also the gut microbiome, relatively little is known about the anti-phage arsenal of Actinobacteria. In this work, we used DefenseFinder to systematically detect 131 anti-phage defense systems in 22,803 fully sequenced prokaryotic genomes, among which 2,253 Actinobacteria of more than 700 species. We show that, like other bacteria, Actinobacteria encode many diverse anti-phage systems that are often encoded on mobile genetic elements. We further demonstrate that most detected defense systems are absent or rarer in Actinobacteria than in other bacteria, while a few rare systems are enriched (notably gp29-gp30 and Wadjet). We characterize the spatial distribution of anti-phage systems on *Streptomyces* chromosomes and show that some defense systems (*e*.*g*. RM systems) tend to be encoded in the core region, while others (*e*.*g*. Lamassu and Wadjet) are enriched towards the extremities. Overall, our results suggest that Actinobacteria might be a source of novel anti-phage systems and provide clues to characterize mechanistic aspects of known anti-phage systems.

## Introduction

The constant threat of viruses has led bacteria to develop a myriad of anti-phage systems. An impressive diversity of anti-phage molecular mechanisms has been characterized in the last five years, but still very little is known about their ecological roles in natural microbial communities. To date, anti-phage defense systems have been mostly studied in a few model organisms, notably *Escherichia coli* or closely related organisms^1–4^. Still, other types of bacteria can harbor very different genomics, metabolisms, and lifestyles. These changing evolutionary pressures could impact the types of defense systems they encode and/or their ecological implications.

Actinobacteria are ubiquitous gram-positive bacteria, which often have high G+C content. Several unique traits distinguish this phylum. First, many Actinobacteria have a complex life cycle, and can for instance form mycelia, sporulate, or undergo morphological differentiation^5–7^. Additionally, certain actinobacterial genera like *Streptomyces* and *Nocardia* have a remarkably rich secondary metabolism and encode for highly diverse reservoirs of bioactive compounds, making them a major source of clinically relevant drugs^8^. Finally, while most bacterial chromosomes are circular, Actinobacteria comprise several species with linear chromosomes, and notably members of the *Streptomyces* genus ^9^. The linearity of their chromosome strongly impacts the genomic structure and organization of *Streptomyces* species. Notably, their chromosome is compartmentalized between a central region, which encompasses most of the core genes of *Streptomyces* and is flanked by two chromosomal arms encoding genes that are generally less conserved ^10,11^.

Despite being a very diverse and widespread bacterial phyla of great economic importance, relatively little is known about the way Actinobacteria defend themselves against phages^12^. Yet, the unique characteristics of certain Actinobacteria might influence their anti-phage arsenal. For instance, the susceptibility to phage infection of actinobacterial species can vary depending on diverse development stages and cell forms^13^. It was also recently discovered that among the bioactive compounds produced by Actinobacteria, some display anti-phage activity ^14,15^, suggesting links between their biosynthetic ability and the defense systems they encode. Finally, the distribution of anti-phage arsenals carried by linear chromosomes has not been described yet.

To determine if the unique characteristics of Actinobacteria confer to their anti-phage defense phylum-specific characteristics, we aimed to characterize both qualitatively and quantitatively the anti-phage arsenal of Actinobacteria and compared it to current knowledge in other common bacterial phyla.

## Methods

### Availability

A list of all genomes used in this analysis and corresponding phylogenetic information is available in Supplementary Table 1. All defense systems and BGCs detected respectively by DefenseFinder ^16^ and antiSMASH ^17^ are available in Supplementary Table 2 and Supplementary Table 3. The results of prophage detection by Virsorter^18^ are available in Supplementary Table 4.

Data and code used for this analysis are available online at https://github.com/mdmparis/Actinobacteria_defense_systems.

### Figures and statistical analysis

Data analyses were done using pandas v 1.3.4 ^19^. Figures were made with python, using modules matplotlib v3.4.3 3 ^20^, seaborn ^21^, and pycirclize. Statistics were computed using python modules scipy.stats ^22^, statsmodels ^23^ and statannotations.Annotator.

### Genome database

22,803 fully sequenced prokaryotic genomes were downloaded in July 2022 from the RefSeq database ^24^. Among them are 2253 genomes of Actinobacteria, accounting for 790 actinobacterial species. All accession numbers and phylogenetic information are available in Supplementary Table 1).

### Detection of defense systems

DefenseFinder v1.0.9 (with DefenseFinder models v1.2.2) ^16^ was used with default parameters to detect anti-phage genes and resulting anti-phage systems in the genomes of the RefSeq database ^16^. This resulted in the detection of 380,556 anti-phage genes, accounting for 161,234 systems. Among these systems, 13,833 were from Actinobacteria. All results of DefenseFinder detection are available in Supplementary Table 2.

### Detection of BGCs

All actinobacterial BGCs were detected by running antiSMASH v.6.0^17^ on all genomes of Actinobacteria of the RefSeq database. In 2253 genomes, 39,164 BGCs were detected. Different types of BGCs were readily annotated by antiSMASH.

### Characterization of the differential abundance of defense systems in Actinobacteria

To determine which anti-phage systems had a differential abundance in Actinobacteria compared to non-Actinobacteria, we built for each system an estimator designed to evaluate if the system is rarer or more abundant in this phylum compared to other bacteria. To do so, we calculated for each type of defense system the difference between its abundance (*i*.*e*. the average number of this system encoded in one genome) in Actinobacteria and its abundance in non-Actinobacteria. Contrary to frequency, the abundance measure takes into account the fact that some genomes encode several copies of the same system. Because the abundances of defense systems are very heterogeneous, we normalized this difference by the general abundance of the system. By doing so, we obtained the following indicator : 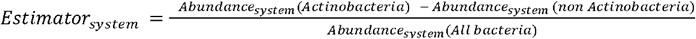

Thus, a negative indicator indicates a system rarer in Actinobacteria than in other bacteria and a positive one indicates a system more abundant in Actinobacteria than in other bacteria (Fig1.C). ANOVA tests corrected by Bonferroni were used to evaluate the difference between the mean number of occurrences of each type of systems per genome encoded by the two groups.

### Building of the phylogenetic trees

For each of the major actinobacterial genera, a core-genome-based phylogenetic tree was generated using PanACoTA v. 1.3.1 ^25^. For each of the genera, we took all the chromosomes (and not plasmids) of the genomes it comprises. For each genus, between one and five chromosomes of closely related genomes were selected as outgroups. We then computed the pan genome of this genus using the PanACotA pangenome module with default parameters (pangenome min identity : 80%), and computed the core genome using the corepers module, with -t 0.95 (meaning that to be considered persistent families need to contain one member in at least 95% of the genomes). Alignments of the core genomes were generated using PanACotA align (default parameters). Finally, the tree module of PanACota was used to generate the corresponding phylogenetic trees using IQtree (default option) with a bootstrap value of 1000. Phylogenetic trees were displayed and annotated using iTOL ^26^.

### Measure of phylogenetic signal on defense systems distribution

Phylogenetic signals were measured for each anti-phage system encoded by *Rhodococcus* and *Streptomyces* were measured using Pagel’s lambda. Both Pagel’s lambda and corresponding p-values were measured using the phylosig function from the phytools R package ^27^.

### Identification of MGE-encoded anti-phage systems and BGCs

The RefSeq annotation of replicon types was used to distinguish plasmids from chromosomes. Among actinobacterial replicons, 2255 were annotated as chromosomes and 878 as plasmids. Virsorter v.2.2.3 (min length set to 2000) was then used to detect prophages in all actinobacterial replicons ^18^. Out of 2253 Actinobacteria genomes, 1656 encoded at least one prophage. The results of Virsorter’s detection are available in Supplementary Table 4.

Due to the possibility of an imperfect detection of prophage boundaries by Virsorter, all defense systems and BGCs that were at least partially included within the boundaries of a given prophage were considered as prophage-encoded. More precisely, multigene anti-phage systems were considered prophage encoded if at least one of their genes was encoded within a prophage, while BGC regions were considered prophage encoded if at least 50% of the region was contained within a prophage.

### Characterization of the spatial distribution of BGCs and defense systems in *Streptomyces*

For the characterization of the distribution of defense systems and BGCS on *Streptomyces* linear replicons, only chromosomes or plasmids annotated as linear were selected. Out of 345 *Streptomyces* chromosomes, 12 were annotated as circular. Although this is most likely misannotations, these chromosomes were excluded from our analysis. Out of 183 *Streptomyces* plasmids, 120 were annotated as linear. As *Streptomyces* are known to encode both linear and circular plasmids, only these 120 plasmids were selected for this analysis.

Each defense system encoded on a linear *Streptomyces* replicon (plasmid or chromosome) was then attributed a relative position, ranging from 0 to 1, by normalizing its start position by the length of the replicon it is encoded on. Similarly, each of the BGCs detected by antiSMASH on a linear plasmid or chromosome was assigned a relative position by normalizing their start position by the size of the replicon that encoded them.

## Results

### Distribution of defense systems in Actinobacteria

To characterize the anti-phage arsenal of Actinobacteria, we used DefenseFinder ^16^ to detect all known anti-phage systems encoded in the genomes of the RefSeq database ^24^ which contains 22,803 fully sequenced genomes (with 2,253 actinobacterial genomes belonging to 790 species). In this dataset, not all actinobacterial species are equally represented, and while three genomes or less are available for 725 of these 790 species, *Mycobacterium tuberculosis* alone represents 267 genomes. In the 2,253 actinobacterial genomes, we detected a total of 161,234 defense systems, among which 13,833 were found in Actinobacteria.

Our analysis reveals that Actinobacteria encode on average 6.14 defense systems per genome (Fig1.A) and dedicate 0.59% of their genomes to anti-phage defense. Although these numbers are slightly lower than the average in bacterial genomes (7.10 systems per genome on average, representing 0.65% of the genome), the most important variation does not appear to be between bacterial phyla, but rather between individuals of a given phylum (Supp Fig1.A). Notably, 36 actinobacterial genomes encode no known defense systems, while the maximum number of defense systems in one actinobacterial genome is 21 (*Olsenella sp*. and *Actinoalloteichus fjordicus*). Out of the 36 genomes with no detected defense systems, 10 belong to species known to have an intracellular lifestyle (*Mycobacterium avium, Mycobacterium intracellulare, Mycobacterium leprae, Mycobacterium lepromatosis*). The same observation has been made in the past for non-Actinobacteria genomes, suggesting that bacterial lifestyle or bacterial phylogeny might influence the anti-phage arsenal of Actinobacteria^16^.

Anti-phage defense systems can be grouped into different types and subtypes, depending on their mechanism and genomic architecture ^28,29^. To characterize the distribution of different types of defense systems in Actinobacteria, we computed their frequencies in this phylum. (Fig 1B, row “All”). Similar to what has been previously described in other bacterial phyla ^16,30^, we observed important heterogeneity in the frequencies of defense systems. While three systems (RM, CRISPR-Cas and Wadjet) are found in more than 40% of actinobacterial genomes, 63 systems are encoded in less than 5% of them (Fig1.B). Interestingly, while the frequency of RM and CRISPR-Cas systems in Actinobacteria (respectively 0.89 and 0.43) is very close to what has previously been reported in all bacteria ^16,30^, the Wadjet systems are a lot more abundant in Actinobacteria than in other bacteria (respectively 42% of Actinobacteria and 7% of non-Actinobacteria encode at least one Wadjet System, Fig1.C) ^2,31^.

**Fig 1:**
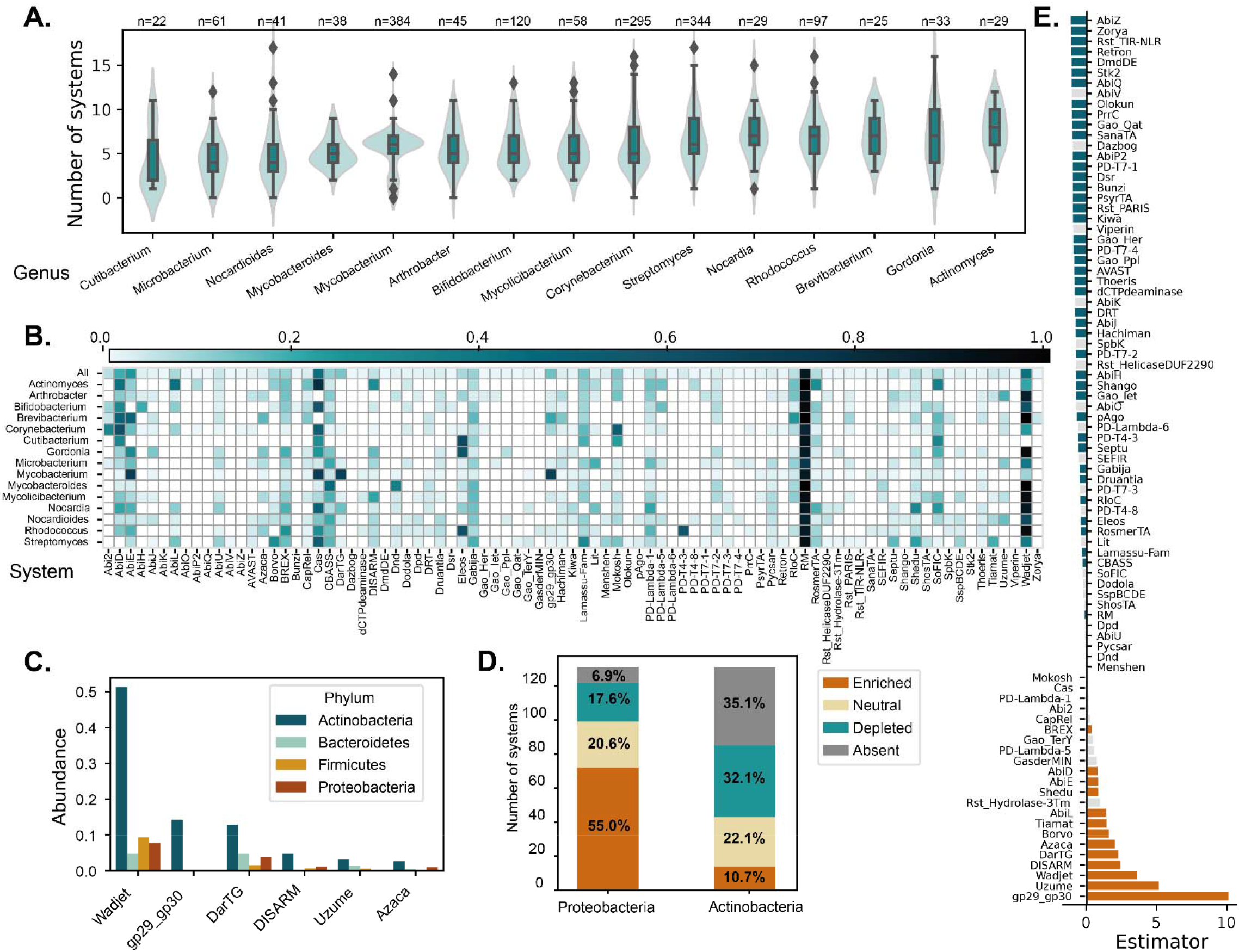
Distribution of defense systems in Actinobateria. **A.** Number of defense systems encoded per genome depending on genera of Actinobacteria. The x-axis iscut at n = 19. **B.** Proportion of the genomes of genera of Actinobacteria that encode different types ofdefense systems. In **A.** and **B.**, only genera containing more than 20 fully sequenced genomes were repre-sented. **C.** Estimators (see formula in Methods) of the differential abundance of different types of actinobac-terial systems compared to non-Actinobacteria. Colored bars (orange: enriched, blue : depleted) representa significant difference of the abundance of a system in Actinobacteria compared to non-Actinobacteria (p ≤ 0.05, ANOVA corrected by Bonferroni). **D.** Proportion of defense systems that are absent, enriched ordepleted compared to other bacteria in Actinobacteria versus in Proteobacteria. **E.** Frequency in majorbacterial phyla of the four types of systems the most enriched in Actinobacteria

Since the frequency of certain systems (*e*.*g* Wadjet) differs in Actinobacteria compared to other bacteria, we then aimed to systematically compare the abundance of the different types of defense systems in Actinobacteria versus in other bacteria. We observed that among the 131 systems that can be detected by DefenseFinder in all prokaryotes, the vast majority are either fully absent (46 systems) from Actinobacteria or are rarer than in other bacteria (42 systems) (Fig1.D and E).

On the other hand, a minority of systems (14 systems) are more abundant in Actinobacteria compared with other bacteria (Fig 1.D and E). The five systems with the highest enrichment score are systems that are rare in other bacterial phyla and one of them (gp29-gp30) is only detected in Actinobacteria (Fig 1.C and Supp Fig 1.C). As a comparison, more than half of the systems encoded by Proteobacteria are more abundant in this phylum than in other bacteria (Fig 1.D and Supp Fig1.B).

Overall, our results suggest that the anti-phage arsenal of Actinobacteria is diverse and currently characterized by a few abundant systems and many rare ones, with important heterogeneity between individuals. The distribution of defense systems appears to be different in Actinobacteria compared to non-Actinobacteria, and some defense systems even appear to be phylum specific.

To better illustrate the diversity of actinobacterial anti-phage arsenals, we represented some examples as chromosome maps in Fig 2.A. Certain Actinobacteria encode clusters of defense systems, with up to 6 distinct defense systems separated by less than 20 proteins. This observation suggests that the tendency of defense systems to colocalize together on the bacterial chromosomes in defense islands ^32^ is also verified in Actinobacteria. Some examples of defense islands are displayed in Fig 2.B.

**Fig 2:**
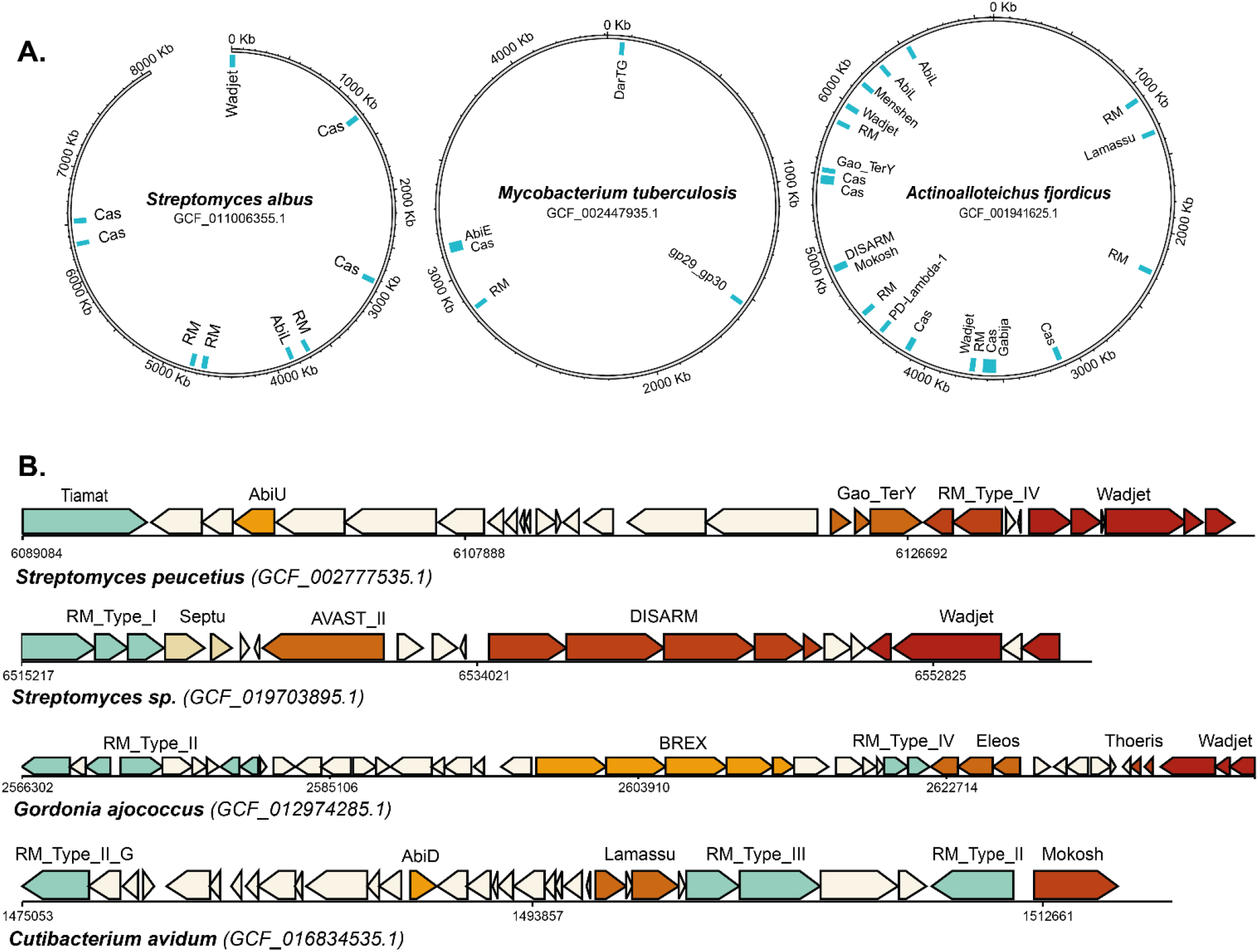
Representatives of the diversity of actinobacterial anti-phage arsenal. **A.** Schematic chromosome maps representing the antiphage arsenals of three Actinobacteria of particular interest: *Actinoalloteichus fjordicus*, one of the Actinobacteria encoding the more defense systems (21 systems); *Mycobacterium tuberculosis*, a major human pathogen; *Streptomyces aibus*, an actinobacterialmodel organism. **B.** Examples of clusters of defense systems found on actinobacterial chromosomes.

### Phylogeny reveals both shared and different systems within members of actinobacterial genera

To assert how the phylogeny of Actinobacteria impacts the anti-phage arsenal they encode at a lower taxonomic level, we first examined the mean number of defense systems encoded in a genome depending on the genus (Fig 1.A). We observed important variations in the number of defense systems encoded between individuals of the same genus, while no major differences could be established between genera of Actinobacteria. When looking this time at the frequency of different systems depending on the actinobacterial genera, we noticed important disparities. For instance, PD-T4-3 and Eleos are extremely enriched in the *Rhodococcus* genus, while CRISPR-Cas systems are completely absent from the genomes of *Brevibacterium* and *Arthrobacter* of the RefSeq database (respectively 25 and 45 genomes) (Fig 1.B).

We then sought to characterize the link between phylogeny and defense systems distribution at the species level. To do so, we built for each of the major genera of Actinobacteria a phylogenetic tree (Fig 3.A and B, Supp Fig 2). As expected from Fig 1.B, some systems are very abundant while others are very rare in a given genus. The most widespread systems in a given genus are often conserved and spread evenly throughout its branches, rather than constrained to only part of the genus (Fig 3.A and B, Supp Fig 2). For instance, in the case of *Rhodoccus (*97 genomes accounting for 17 species), 73% of the genomes encode a Wadjet and a RM system and either a Eleos or a PD-T4-3 system (Fig 3.A). Similarly, in the case of *Streptomyces* (344 genomes representing 141 species), 92% of the genomes encode at least one RM system and 70% of them encode at least two RM systems (Fig 3.B). On the other hand, certain systems are very rare and unevenly distributed between members of a given genus. For instance, in the *Streptomyces* genus, 28 systems are encoded by less than 5% of the genomes and are disseminated evenly throughout the tree, while in *Rhodococcus*, 22 systems are found in less than 5% of the genomes (Fig 3.A, Fig 3.B). To determine whether these rare systems were scattered throughout the tree or confined to specific subclades, we computed for each of *Rhodococcus* and *Streptomyces* systems a measure of the phylogenetic signal with Pagel’s λ (see Methods). For most of the systems encoded by less than 5% of *Streptomyces* and *Rhodococcus* genomes (respectively 23 and 17 systems), we observed no significant phylogenetic signal (p value > 0.95), meaning that pairs of closely related strains were no more similar to each other than two randomly selected strains. This suggests that most rare systems in these genera have a patchy distribution.

**Fig 3:**
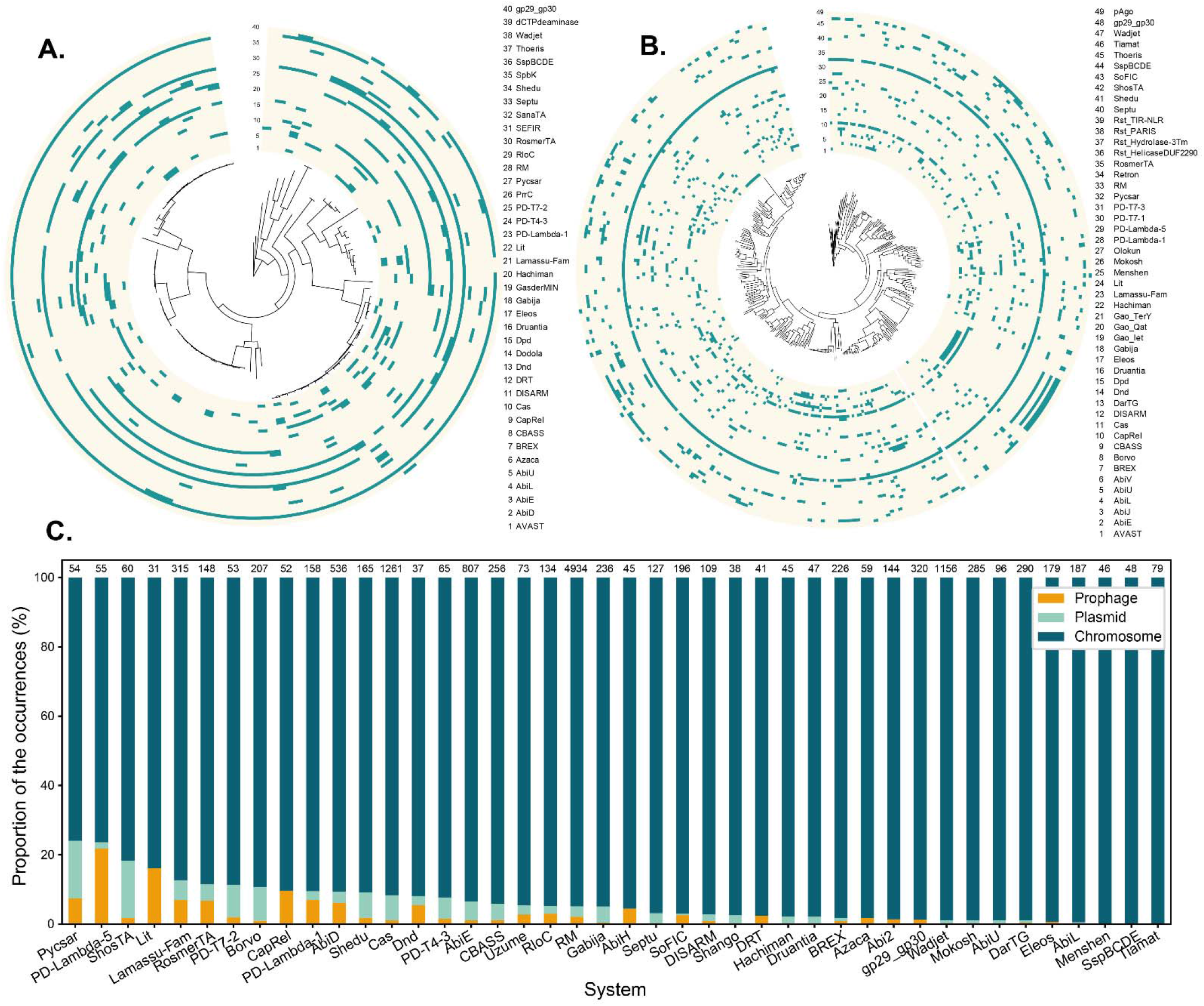
Evidences of Horizontal Gene Transfer in actinobacterial defense systems. **A and B:** Presence (blue) or absence (light yellow) of different types of defense systems in the genomesmapped on the phylogenetic trees (generated using PanACoTA based on the core genome) of two major actinobacterial genera, respectively *Rhodococcus* **(A.)** and *Streptomyces* **(B.)**. Both trees are rooted on outgroups tothe genus **C.** Relative contribution (in % of the occurrences) of different types of genetic elements to differenttypes of defense systems. Numbers above each bar indicate the total number of occurences of each type ofsystems.

Overall, the different genera of Actinobacteria seem to often encode a combination of systems that are very widespread within its members and of very rare systems with a patchy distribution. Both the types of shared and variable defense systems vary depending on the genus.

### MGEs contribute to the anti-phage arsenal of Actinobacteria

Mobile Genetic Elements (MGEs), such as plasmids, integrative and conjugative elements and phages are known to carry defense systems and contribute to the fast exchange rate of defense systems in bacteria ^3,4,33–36^. Because the contribution of MGEs to the anti-phage arsenal of bacteria has rarely been characterized outside *Proteobacteria*, we tried to determine this phenomenon in Actinobacteria. To do so, we restricted our analysis to two types of MGEs, plasmids, and prophages. We determined for each defense system detected if it is encoded on a plasmid, or within a prophage. Out of the 13833 defense systems detected in Actinobacteria by DefenseFinder, 2.0% (279) are encoded within a prophage, and 3.4% are encoded on a plasmid (469).

Since we observed a heterogeneous distribution of defense systems within Actinobacteria, we then aimed at determining whether some systems are more frequently encoded by MGEs than others. To do so, we calculated for each type of defense system the proportion of the occurrences of the system in a phage, in a plasmid or simply in the chromosome (Fig 3.C). We observed that the proportion of systems encoded on an MGE varies depending on the type of system considered. Some systems are almost always chromosomal (*e*.*g*. like Tiamat, or Ssp), while MGEs are important contributors to some other systems. Among the systems that are often encoded by MGEs, some tend to be plasmid-encoded, like ShosTA, while others tend to be prophage-encoded, like PD-Lambda-5 and Lit. The two systems that are the most frequently found on prophages were both discovered in *E*.*coli* prophages (e14 prophage for the Lit system ^37^, and P2-like prophage for PD-Lambda-5 ^3^, suggesting that the propensity of certain systems to be encoded by prophages might be conserved across bacterial phyla.

Because we had previously observed that the distribution of defense systems is highly variable within Actinobacteria genera, we wondered if the propensity to be MGE-encoded of a given system is linked to its abundance in a given bacterial clade. For each of the major genera of Actinobacteria, we looked at the proportion of occurrences of each type of system that is encoded on an MGE (plasmid or prophage), depending on the proportion of genomes that carry this type of system in the genus (Supp Fig 3.A). We observed that the systems encoded in more than 50% of the genomes of a given genus were rarely encoded on phages and plasmids.

A lower GC percentage compared to the rest of the genome is a known marker of HGT in bacteria. To explore the link between HGT and the abundance of a system in Actinobacteria, we computed a GC score for each anti-phage gene by normalizing its GC percentage by the GC percentage of the replicon it is encoded on. Thus, a GC score below 1 means that the gene is enriched in AT compared to the rest of the replicon. We then looked at the average GC ratio of each type of system depending on the proportion of genomes that carry this type of system in a given genus (Supp Fig 3.B) We observed that in all genera studied, the systems with a GC ratio below 0.8 were generally rare systems. These observations suggest that the rarest systems in a given genus are also the ones that are the most frequently exchanged through HGT, at least partly due to MGEs.

Therefore, it appears that MGEs can be associated with defense systems in Actinobacteria, as observed for other bacterial phyla and notably Proteobacteria. This association probably supports HGT of defense systems between bacteria. MGEs do not contribute equally to all types of defense systems. It also seems that the contribution of MGE to anti-phage arsenals varies depending on the genus, which does not seem linked to differences in the number of MGEs encoded by genomes of the genus (Supp Fig 4.A and B).

**Fig 4:**
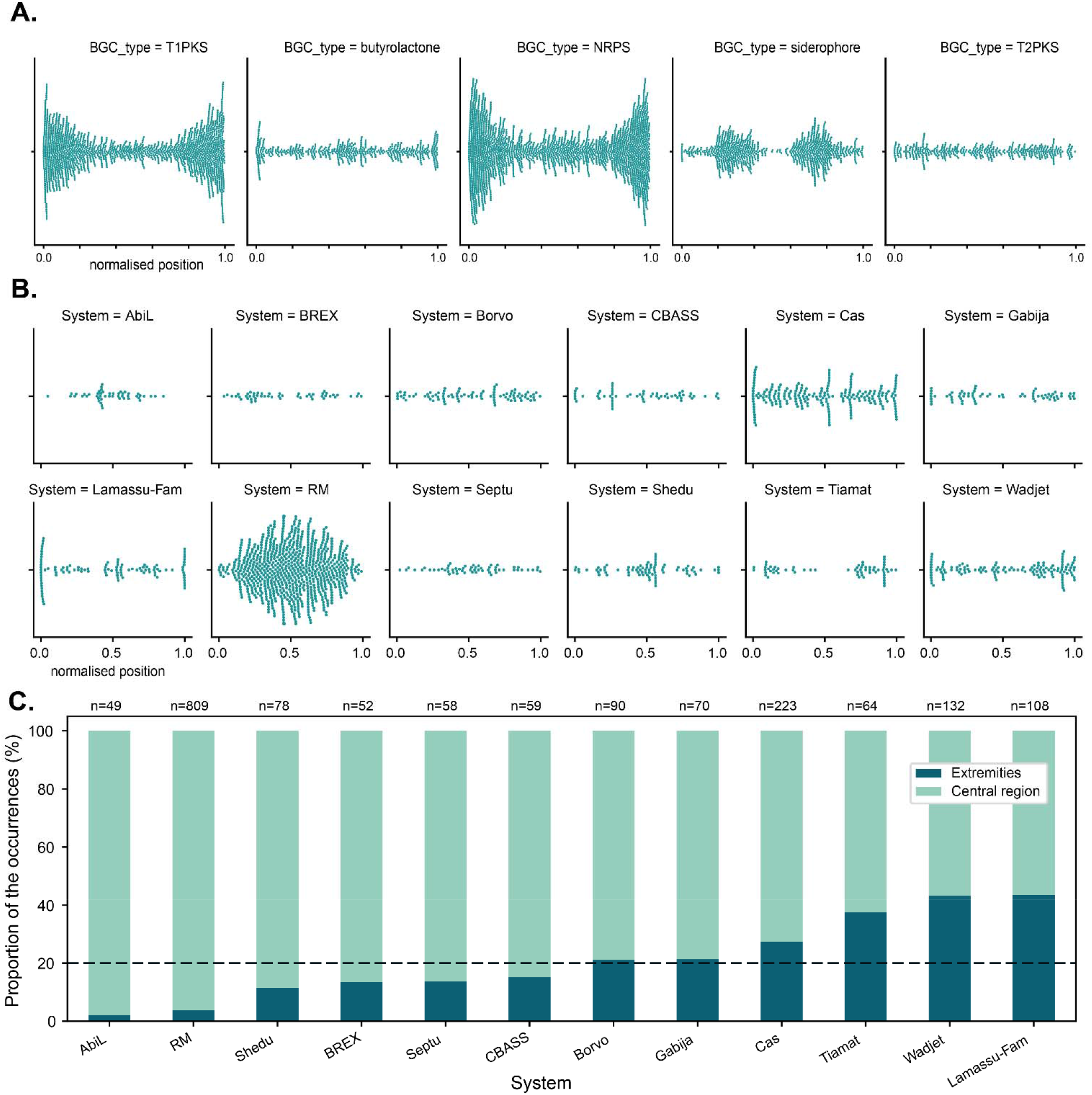
Patterns of spatial distribution of defense systems along Streptomyces chromosomes revealsparallels with BGCs. **A.** Archetypal examples of the distribution of the normalised position of BGCs along *Streptomyces* linear chromosomes. **B.** Distribution of the normalised position of defense systems on *Streptomyces* linear chromosomes. **C.** Proportion of each types of defense systems that are encoded in the extremities (first or last 10%) of the chromosome, versus the ones encoded in the middle (80%) of the chromosome. The dotted black line indicates theproportion of the chromosome represented by the extremities as defined here, *i.e.* 20%. Above each bar isindicated the total number of occurrences of each type of systems in *Streptomyces* linear chromosomes. In **B.** and **C.**, only systems with more than 45 occurrences are represented.

### Comparison of the spatial distribution of defense systems and BGCs reveals conserved trends between two types of gene clusters involved in biological interactions

Actinobacteria are known to carry numerous Biosynthetic Gene Clusters (BGCs), which produce a striking diversity of secondary metabolites ^6,38^. Many of these metabolites are bioactive molecules involved in biological interactions ^39^, for instance with other bacteria or with fungi. It was recently discovered that several secondary metabolites produced by Actinobacteria had a strong anti-phage activity, a phenomenon that was termed anti-phage chemical defense ^14,15^. This discovery highlights a parallel between BGCs and anti-phage systems in Actinobacteria. Indeed, both are locally adaptive systems that can be used in biological interactions. Because of this parallel, we sought to compare BGCs and defense systems to see if they present similar patterns.

To do so, we used antiSMASH v.6 ^17^, to systematically detect all known BGCs in the Actinobacteria genomes of the RefSeq Database. We found that on average, an Actinobacteria genome encodes 17.4 BGCs. Contrary to what we observed with anti-phage systems, the number of BGCs per genome is highly variable depending on the genus (Supp Fig 5.A). As expected from the literature ^38^, the *Nocardia* and *Streptomyces* genera are much more BGC-rich than other genera, with on average respectively 45 and 46 BGCs per genome. Like for defense systems, we also observe an important heterogeneity in the frequency of different types of BGCs detected by antiSMASH depending on the genus (Supp Fig 5.B). Additionally, we found that BGCs were often plasmid-encoded, and a few seemed to be encoded in prophages (Supp Fig 5.C). These observations are in line with the multiple reports of the importance of HGT in BGCs evolutionary dynamics in Actinobacteria ^40,41^, similarly to defense systems.

Not only are certain actinobacterial genera characterized by a rich secondary metabolism, but some, and in particular *Streptomyces*, also have linear chromosomes. *Streptomyces* linear chromosomes are spatially compartmentalized, with a relatively conserved central region and more divergent chromosomal arms. Several studies show that the arms of *Streptomyces* chromosomes are enriched in BGCs ^10,42,43^, but the position of defense systems on these chromosomes has not been characterized. To investigate the consequences of the linearity and compartmentalization of the *Streptomyces* chromosome on anti-phage defense and compare it to what has been observed for BGCs, we selected the chromosomes of all fully sequenced *Streptomyces* genomes available (344 genomes of 141 species). The length of these chromosomes was highly variable (ranging from 1,9 Mbp to 12,3 Mbp with a median of 8,1 Mbp). A recent study on more than a hundred *Streptomyces* chromosomes shows that both the length of the chromosomal arms and of the core region correlate with genome size ^44^. Thus, we used a normalized position to systematically characterize the spatial distribution of defense systems and compare it with BGCs.

As a baseline to characterize the distribution of genes involved in biological interactions, we first mapped the distribution of BGCs along *Streptomyces* chromosomes. As expected from the literature, we observed that certain BGC types (*e*.*g*. Type 1 PKS, Type 3 PKS, NRPS⋯) are strongly enriched towards the extremities of the chromosome (Fig4.A and Supp Fig 6.A and B). However, we also observe that certain BGCs do not appear to be specifically encoded in the chromosomal arms but are homogeneously distributed along the chromosome (T2PKS for instance). Mapping this time the distribution of defense systems, we also observed specific patterns of spatial distribution that echoed the ones observed for BGCs (Fig 4.B). On one hand, some systems appear to be mostly encoded toward the center of the chromosome, with the most striking example being RM systems. On the other hand, some systems, such as Lamassu and Wadjet systems, are mostly encoded towards the extremities of the chromosome. As it was previously reported that the length of the arms correlates with the length of the chromosome in *Streptomyces* ^44^, we calculated the proportion of the occurrence of each defense system within these arms, represented here as the first or last 10% of the chromosome (Fig 4.C). Less than 4% of the RM systems are encoded in the chromosomal arms, while this proportion reaches more than 43% for Lamassu and Wadjet systems. This observation suggests at least two types of spatial organization. We also noticed that Lamassu systems are particularly enriched at the very ends of the chromosome, as almost a third (31%) of Lamassu systems are found in the first or last 1% of the chromosome (1% of the chromosome representing on average 83 kb in this case) (Supp Fig 7.A). Strikingly, Lamassu systems encoded in the first or last 1% of the chromosome are evenly distributed around the *Streptomyces* phylogenetic tree (Supp Fig 7.B). *Streptomyces* often encode linear plasmids that share their general structural organization with linear chromosomes ^9^. The Lamassu systems encoded on one of the 120 *Streptomyces* plasmids annotated as linear on RefSeq also tended to be encoded near the extremities of linear plasmids (Supp Fig 7.C).

Overall, anti-phage defense presents unique spatial distribution patterns in *Streptomyces*, which are linked to the unique genomic organization of these bacteria. These patterns echo the ones observed for BGCs, suggesting that they could be characteristic not of anti-phage defense, but rather of genes involved in biological interactions.

## Discussion

### Discovery bias: more systems to discover in Actinobacteria and their MGE

In this work, we propose a comprehensive view of the anti-phage arsenal of Actinobacteria. We first observe that Actinobacteria encode many systems. Strikingly, approximately a third of the systems that have been described to date were not detected at all in Actinobacteria. Additionally, among systems encoded by Actinobacteria, most were rarer in this phylum than in other bacteria. One possible explanation for the absence or reduced abundance of most known systems in Actinobacteria could be that they encode systems that are different from commonly characterized bacteria and that have not yet been discovered. Supporting this hypothesis, both the bioinformatic and experimental discovery methods of anti-phage systems are strongly biased toward certain phylogenetic groups, and notably the Proteobacteria phylum ^2–4^. Out of the 131 systems detected by DefenseFinder, only three systems were either from or tested in Actinobacteria (Ssp, Pgl and gp29-gp30), against 99 systems from or tested in Proteobacteria. Coincidentally, most defense systems are enriched in Proteobacteria, while only a few are enriched in Actinobacteria. This observation reinforces the idea of a discovery bias, favoring notably *E. coli* and closely related organisms.

Efforts to systematically discover prophage-encoded systems are also strongly biased toward Proteobacteria ^4,33^. Indeed, we found that the 6 defense systems that were previously reported as the most prophage-encoded in all prokaryotic genomes (above 60% of the total number of systems) ^16^ were almost exclusively found in Proteobacteria (99% of the occurrences of these 6 systems). A handful of systems were recently discovered in *Mycobacterium* phages ^45–48^, which are by far the most characterized and studied actinobacterial phages ^49^. Only one of these systems, gp29-gp30, is currently detected by DefenseFinder, and it is exclusively found in Actinobacteria. This supports the idea that novel phylum-specific anti-phage systems can be discovered in bacteriophages infecting Actinobacteria.

Recently, studies focused on Actinobacteria have uncovered novel anti-phage strategies, including transient loss of the cell wall ^50^ and production of anti-phage compounds ^14,15^. Our results suggest that many more anti-phage systems might be discovered in Actinobacteria and their MGEs.

### Comparison between BGCs and defense systems reveals both shared and distinct characteristics

Because BGCs and anti-phage systems are both conditionally adaptive gene clusters that are often involved in biologic conflicts, we compared these two types of systems in Actinobacteria. We found that both are abundant and widespread in this phylum. Like defense systems, BGCs can be carried by MGEs (Supp Fig 6.C), which probably participates in their HGT.

In the linear chromosomes of the genus *Streptomyces*, we observe that both defense systems and BGCs have two types of distribution. Because *Streptomyces* chromosomes are known to be spatially compartmentalized between a core and a variable region, we initially expected these distributions to be linked to the abundance of the systems (either BGCs or anti-phage). However, the propensity of the relatively abundant Wadjet and Lamassu systems to be encoded in the extremities of the chromosomes seems to contradict this idea. Other hypotheses could explain the observed patterns of distribution along *Streptomyces* chromosomes. Notably, and since the level of gene expression is known to vary depending on the position on the chromosome, the spatial distribution of defense systems and BGCs could reflect different types of regulation. The different patterns of spatial distribution could also be caused by different types of ecological roles or evolutionary dynamics.

### *Streptomyces* linear chromosomes as a model to explore molecular mechanisms of defense systems

The spatial distribution of anti-phage systems on *Streptomyces* linear chromosomes could also provide hints on their mechanisms. Strikingly, the two systems that are the most frequently found at the extremities of the chromosomes (Lamassu and Wadjet) both encompass SMC domain, have anti-plasmid activity, and have been proposed to sense the structure of plasmid DNA ^31,51,52^. This suggests that the spatial distribution of defense systems might have links with their molecular mechanisms.

Additionally, Lamassus seem to be encoded at the very extremities of the arms. Streptomyces are known to encompass Terminal Inverted Repeat sequences (TIRs) flanking the chromosomal arms at each extremity of the chromosome and of the linear plasmids ^9,53^. The size of these regions varies both intra- and interspecifically, with sizes ranging from less than 1kb to up to 1Mb for *S. coelicolor* A3(2) ^53,54^. It is plausible that some of the Lamassus systems that we found to be encoded at the very extremities are part of the TIR region of the chromosome or plasmid. The conservation of Lamassu systems in a region known for its variability suggests a possible shared biological function. Exploring patterns of defense systems spatial organization more in depth might provide insights on the function of anti-phage defense systems that remain elusive.

Overall, our results highlight the importance of studying defense systems outside of commonly studied organisms (*e*.*g. B. subtilis* and *E. coli*), not only to discover novel anti-phage systems but also to characterize the ones that are currently known, both from a mechanistic and ecological point of view.

## Acknowledgments

We would like to thank Julia Frunzke, Dennis Claessen, Véronique Ongenae and Daniel Rozen for fruitful discussions and feedback on this work. We are also grateful to members of the MDM lab for their very useful comments on earlier versions of the manuscript.

