## Supplementary Figures for "Genomic characterization of the antiviral arsenal of Actinobacteria"

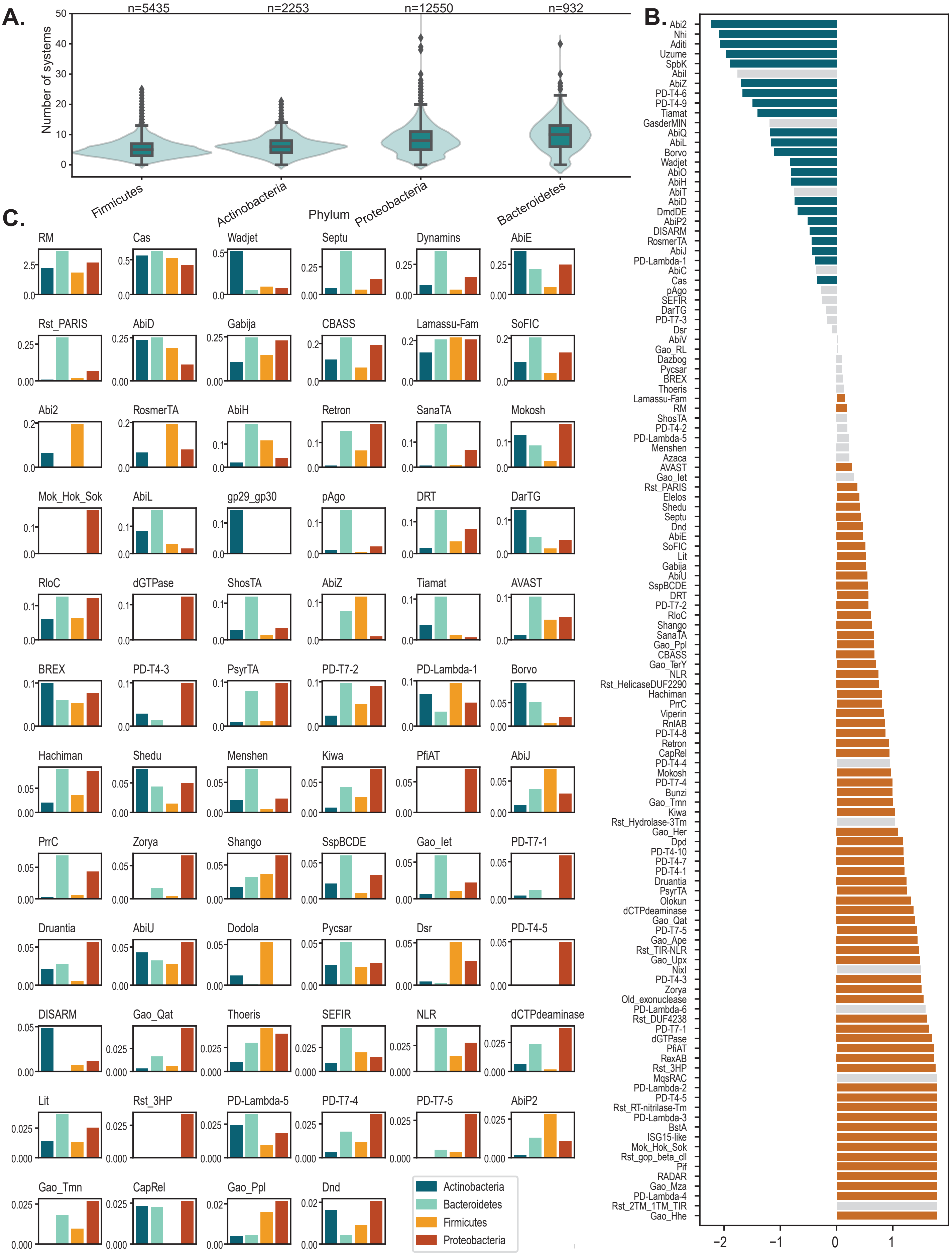

**Supp Fig 1: Distribution of defense systems depending on the bacterial phyla**

**A.** Number of systems per genome in 4 major bacterial phyla (> 500 genomes per phylum). Y-axis was cut at 50, the maximum number of systems in one genome being 64. **B.** Enrichment score of defense systems of Proteobacteria vs non-Proteobacteria. Colored bars (orange: enriched, blue : depleted) represent a significant difference of the abundance of a system in Actinobacteria compared to non-Actinobacteria (p≤0.05 , ANOVA corrected by Bonferroni). **C.** Abundance of different defense systems in 4 bacterial phyla. Y-axis represents the average number of a given type of system in one genome. Only systems with more than 300 occurrences in all prokaryotic genomes are represented

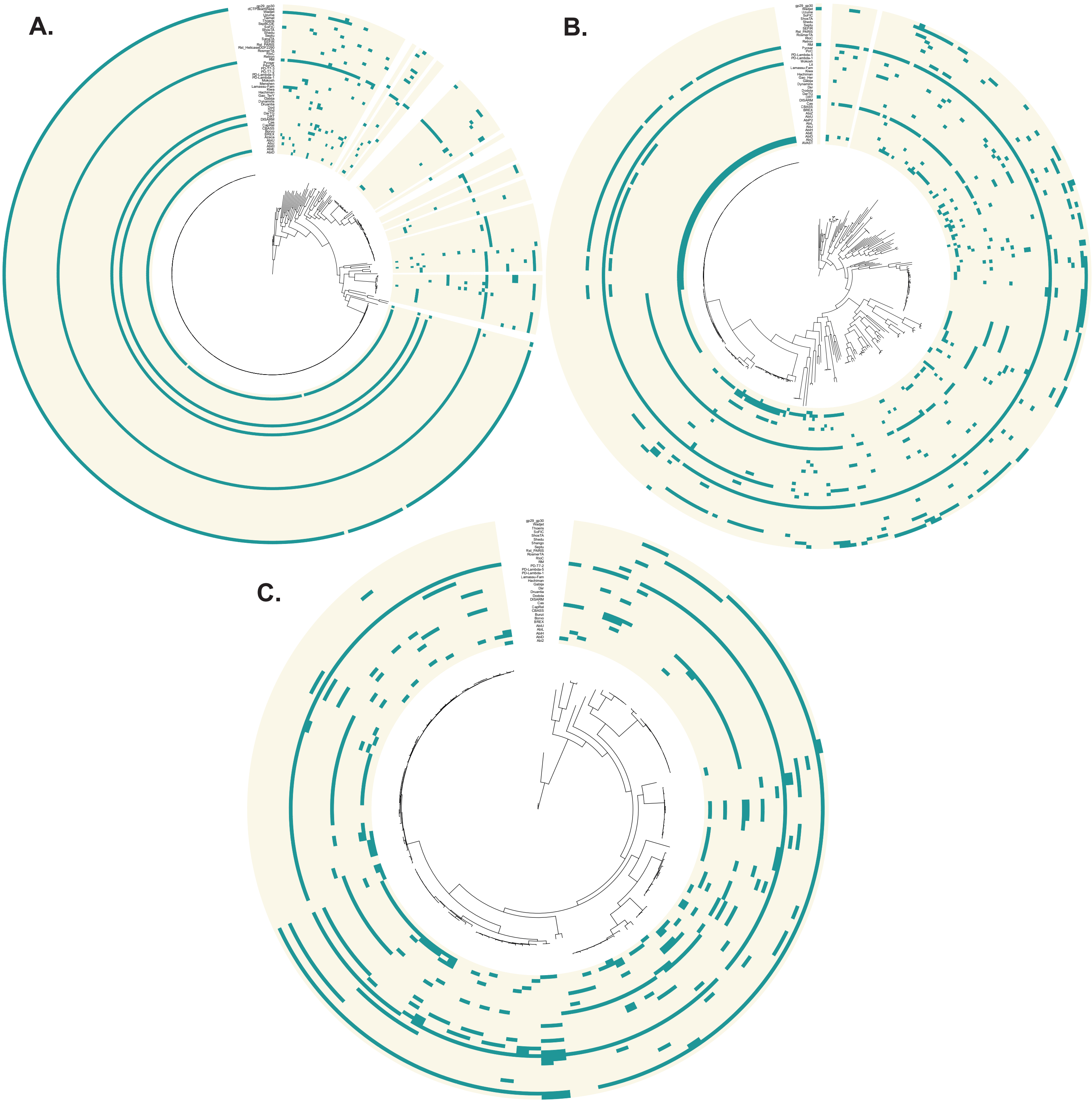


**Supp Fig 2 : Distribution of defense systems in three major actinobacterial genera**
Presence (blue) or absence (light yellow) of different type of defense systems in the genomes mapped on the phylogenetic trees (generated using PanACoTA based on the core genome) of three major actinobacterial genera, respectively *Mycobacterium* **(A.)**, *Corynebacterium* **(B.),** and *Bifidobacterium* **(C.)**. All trees are rooted on outgroups of the genus.

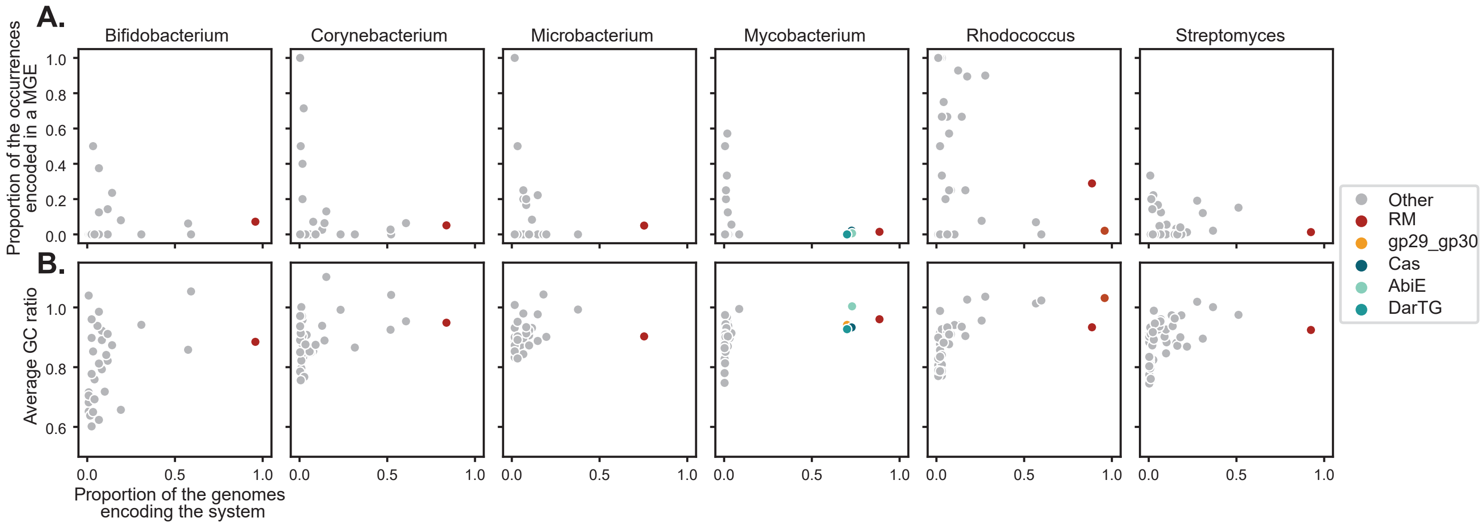


**Supp Fig 3: Distribution of defense systems depending on the bacterial phyla**

In A. and B. only genera containing more than 60 genomes are represented. **A.** Proportion of the occurrences of each given type of systems that are encoded on a plasmid or a prophage depending on the proportion of the genomes that encode this type of system. **B.** Average GC ratio (= GC of the system / GC of the replicon) of different types of defense systems depending on the proportion of the genomes that encode this type of system.

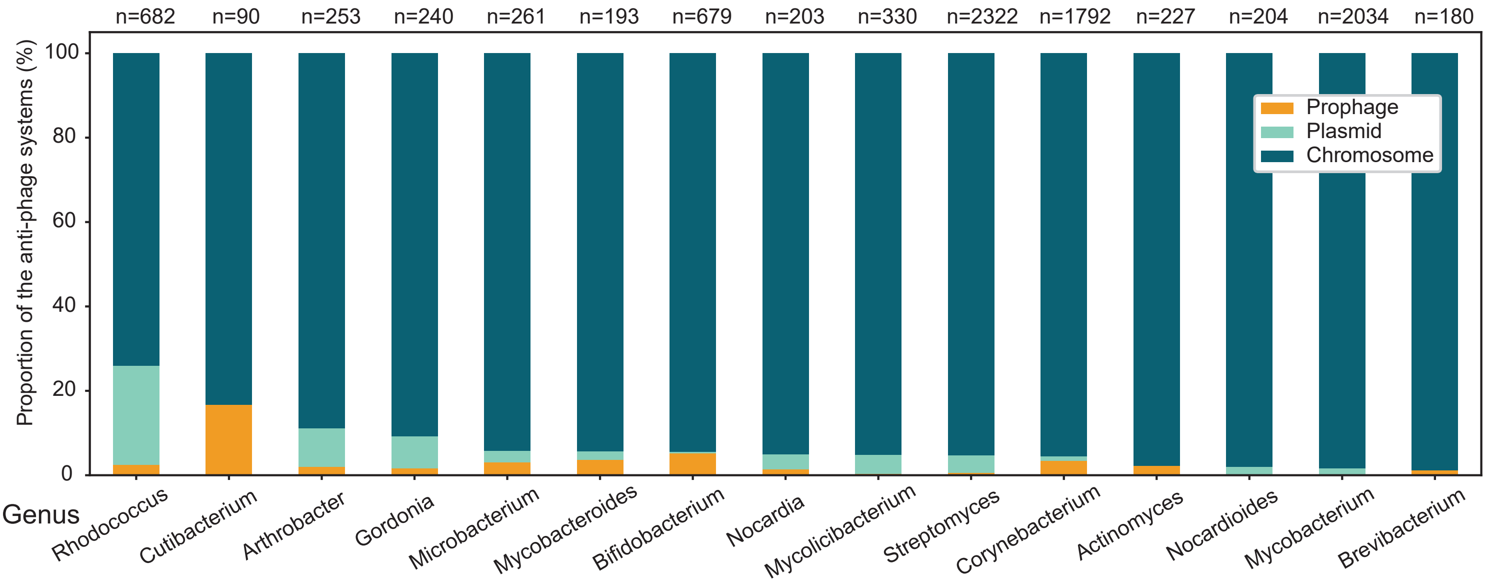


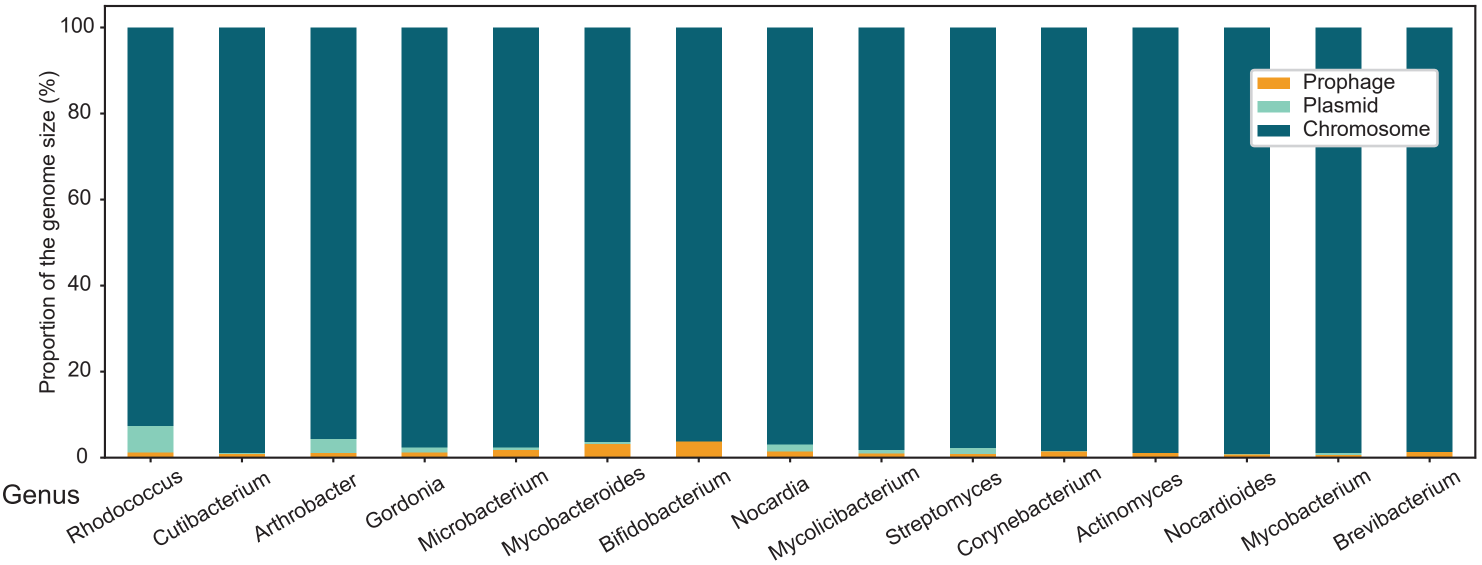


**Supp Fig 4 : Differential contribution of MGEs to the anti-phage arsenals of different actinobacterial genera**

**A.** Proportion of the defense systems of a given genus that are encoded by different types of genetic elements (chromosomes, plasmids, phages). Above each bar is indicated the total number of systems in the genus. **B.** Relative contribution of MGEs to the total genome length of different genera. Color bars represent the proportion of the total genome length of all genomes in a given genus represented by different types of genetic elements (chromosomes, plasmids, phages).

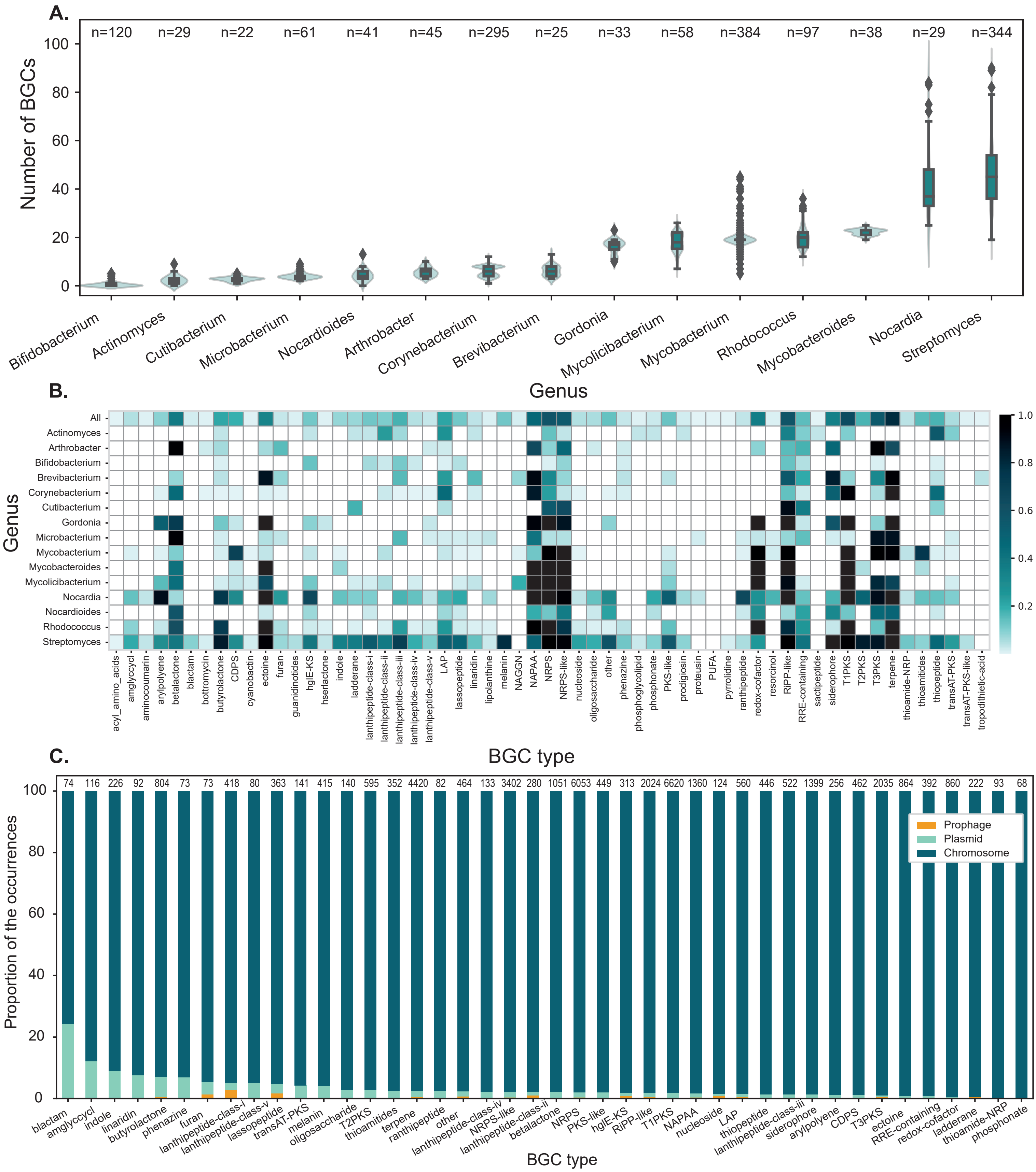


**Supp Fig 5 : Distribution of BGCs in Actinobacteria**

**A.** Number of BGC per genome in different genera of Actinobacteria. **B.** Frequency of different types of BGC in different genera of Actinobacteria. In **A.** and **B.,** only genera containing more than 20 genomes are represented. **C.** Relative contribution of MGEs to different types of BGCs. Color bars represent the proportion of BGCs of a given type that are encoded by different types of genetic elements (plasmids, phage, chromosome). Only BGC types with more than 60 occurrences are represented.

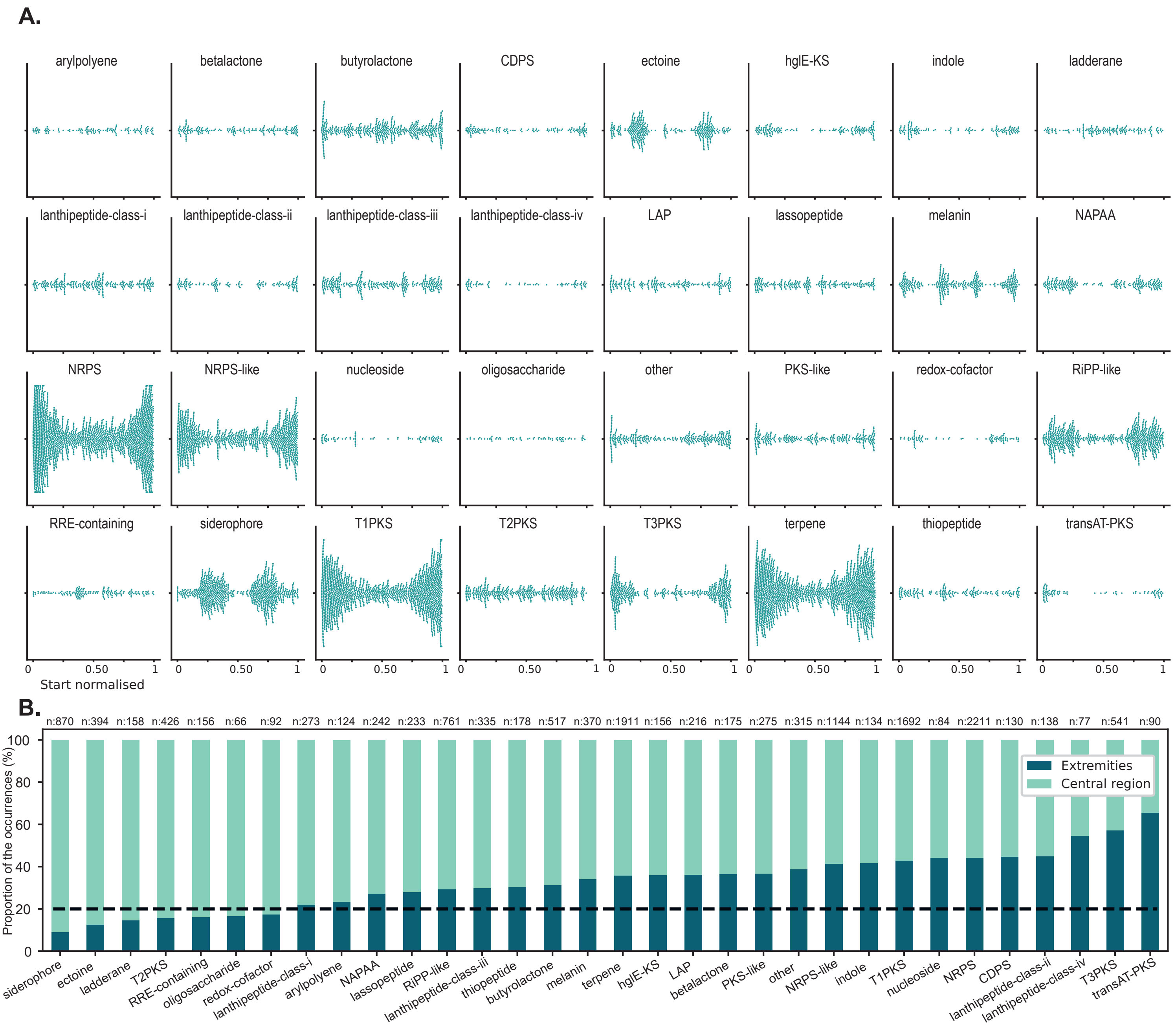


**Supp Fig 6 : Spatial distribution of BGCs along *Streptomyces* chromosomes**
**A.** Distribution of the normalised position of different types BGCs on *Streptomyces* chromosomes. **B.** Proportion of the occurrences of each types of BGCs that are encoded in the central region versus the extremities (first or last 10%) of the chromosome. The number of occurrences of each type of BGCs is indicated above the bars. In **A.** and **B.**, only systems with more than 60 occurrences on linear chromosomes are represented

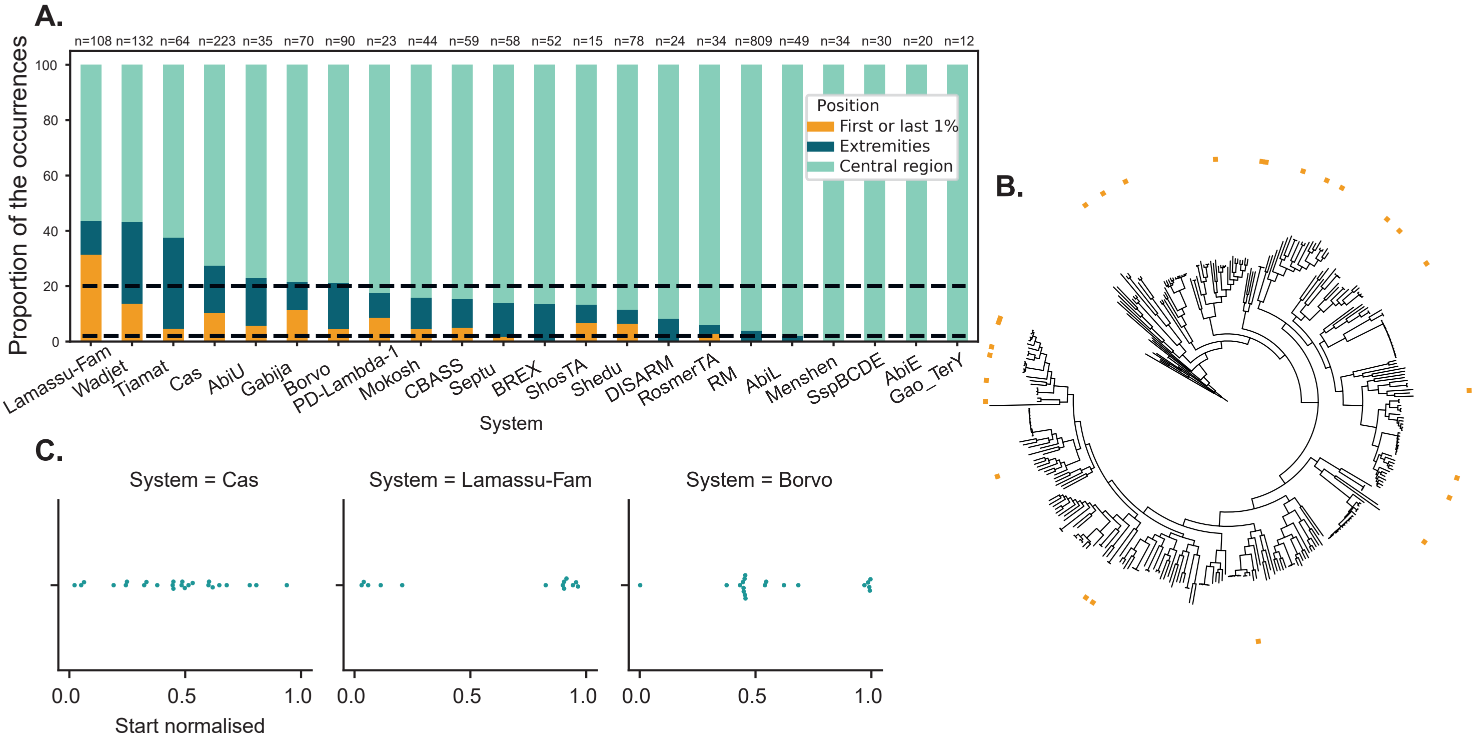


**Supp Fig 7: Lamassu systems are often encoded at the very ends of the chromosome in *Streptomyces***
**A.** Proportion of different defense systems that are encoded in the central region (light blue), in the extremities (first and last 10%, dark blue), or in the very ends of the extremities (first and last 1% of the chromosome, orange). The dotted black lines indicated the proportion of the chromosome represented by each region (20 % for dark blue region and 2% for the orange region). Indicated above each bar is the number of occurences of each type of system. **B.** Distribution of Lamassu systems encoded in the first or last 1% of the chromosome (orange squares) on the phylogenetic tree of *Streptomyces*. **C.** Spatial distribution of defense systems on *Streptomyces* linear plasmids. Only systems with more than 10 occurrences on linear plasmids are represented.
